# Hybrid immunity elicits potent cross-variant ADCC against SARS-CoV-2 through a combination of anti-S1 and S2 antibodies

**DOI:** 10.1101/2023.03.09.531709

**Authors:** Michael D. Grant, Kirsten Bentley, Ceri A. Fielding, Keeley M. Hatfield, Danielle P. Ings, Debbie Harnum, Eddie Wang, Richard Stanton, Kayla A. Holder

## Abstract

Antibodies capable of neutralising SARS-CoV-2 have been well studied, but the Fc receptor-dependent antibody activities that also significantly impact the course of infection have not been studied in such depth. SARS-CoV-2 infection induces antibody-dependent NK cell responses targeting multiple antigens, however, as most vaccines induce only anti-spike antibodies, we investigated spike-specific antibody-dependent cellular cytotoxicity (ADCC). Vaccination produced antibodies that only weakly induced ADCC, however, antibodies from individuals who were infected prior to vaccination (‘hybrid’ immunity) elicited much stronger anti-spike ADCC. Quantitative and qualitative aspects of humoral immunity contributed to this capability, with infection skewing IgG antibody production towards S2, vaccination skewing towards S1 and hybrid immunity evoking strong responses against both domains. The capacity for hybrid immunity to provide superior spike-directed ADCC was associated with selectively increased antibody responses against epitopes within both S1 and S2. Antibodies targeting both spike domains were important for strong antibody-dependent NK cell activation, with three regions of antibody reactivity outside the receptor-binding domain (RBD) corresponding with potent anti-spike ADCC. Consequently, ADCC induced by hybrid immunity with ancestral antigen was conserved against variants containing neutralisation escape mutations in the RBD [Delta and Omicron (BA.1)]. Induction of antibodies recognizing a broad range of spike epitopes and eliciting strong and durable ADCC may partially explain why hybrid immunity provides superior protection against infection and disease than vaccination alone, and demonstrates that spike-only subunit vaccines would benefit from strategies to induce a combination of S1- and S2-specific antibody responses.

**Significance:** Neutralising antibodies prevent the entry of cell-free virus, however, antibodies that promote Fc-dependent activities such as ADCC are critical to control cell-associated virus. Although current SARS-CoV-2 vaccines induce potent neutralising antibodies, they fail to induce robust ADCC. Our demonstration that hybrid immunity induces superior ADCC with pan-variant activity may partially explain why hybrid immunity offers enhanced protection against reinfection. It also highlights that vaccine strategies based on expression of the spike subunit alone should not focus solely on inducing antibody responses targeting the receptor binding domain.

## Introduction

Humoral immunity against SARS-CoV-2 spike (S) protein is induced by infection and by vaccination with any of the predominant COVID-19 vaccines used worldwide, most of which encode S as a single antigen (1–3). Anti-S antibodies (Ab) target multiple regions within the protein, but the major focus has been on those that neutralise cell-free virions. These primarily bind within the receptor binding domain (RBD) or in some cases the N-terminal domain (NTD), both of which are found in the S1 domain of the protein. Neutralising Abs block or prevent binding between SARS-CoV-2 and the entry receptor angiotensin converting enzyme-2 (ACE-2) or prevent post-binding events required for virus entry (4, 5). They are thought to be crucial for reducing transmission of SARS-CoV-2, and thus, are a key measure for predicting COVID-19 vaccine efficacy (6).

Despite their clear importance, neutralising Abs have recognised limitations. The number of neutralising epitopes is limited, resulting in rapid selection of SARS-CoV-2 variants with mutations that weaken Ab binding to key neutralising sites (7, 8). After approximately 3 years of evolution in the human population, SARS-CoV-2 variants of concern have largely escaped the neutralising activity of Abs induced by the ancestral S antigen and are continually evolving to evade Abs induced by infection with more recent variants. As a result, the efficacy of vaccines at preventing infection is already reduced within months of their introduction. Once infection occurs, SARS-CoV-2 can undergo direct cell-cell transmission, which further undermines the efficacy of neutralising Ab (9).

To counteract cell-cell virus spread, Abs are required which, rather than neutralising cell-free virions, recognise viral antigens on the surface of infected cells (10). These recruit effector cells such as natural killer (NK) cells to kill infected cells through antibody-dependent cellular cytotoxicity (ADCC), thereby controlling cell-associated virus. Infection with SARS-CoV-2 readily induces Ab capable of supporting ADCC (11) and ADCC is a key determinant of immunological control in animal challenge models (12–21). We and others have shown that Ab capable of ADCC are effective at preventing disease in animals even in the complete absence of neutralising activity (18, 22). Thus, inducing and maximising this activity through vaccination is highly desirable.

In addition to being a target for neutralising Abs, SARS-CoV-2 S is also expressed on the infected cell surface, where it is efficiently bound by Abs (11). Thus, S has the potential to be an effective target for Fc receptor (FcR)-mediated NK cell activation, providing a single vaccine antigen capable of inducing Ab activity targeting both cell-free and cell-associated virus. Although studies have shown that vaccination induces anti-S Abs capable of activating NK cells when tested against purified protein (23–25) or transfected cells (26–31), our previous study revealed that, when tested against live virus, effective ADCC was dominated by Abs targeting nucleocapsid (N), membrane (M), and ORF3a and individuals who had only anti-S Abs (i.e. vaccinees with no infection history) demonstrated weak ADCC (11).

Subsequent studies have made it clear that infection with SARS-CoV-2 prior to vaccination with a S-encoding vaccine (‘hybrid’ immunity) offers superior protection compared to vaccination or infection alone (32–35). To explore Fc-dependent mechanisms that may contribute to this protection, we examined quantitative and qualitative aspects of S-induced humoral immunity and its association with ADCC potency in individuals who had recovered from infection, who had been vaccinated, or both.

## Results

### Hybrid immunity elicits robust ADCC against S-transduced cells

As all donors in this study were vaccinated with SARS-CoV-2 S-encoding vaccines, we isolated S-specific ADCC from responses targeting other SARS-CoV-2 proteins by testing Abs for their capacity to mediate lysis of SARS-CoV-2 S-expressing target cells. A series of plasma dilutions were performed initially to establish optimal levels on a subset of samples and plasma was diluted 1:1000 for subsequent experiments. This dilution distinguished ADCC from no ADCC for vaccinee and post-infection samples and discriminated ADCC levels from hybrid immunity across a wide range without plateauing.

There was minimal background ADCC against parental (non-transduced) MRC-5 elicited by SARS-CoV-2 seropositive plasma and against Wuhan-Hu-1 S-expressing MRC-5 (Wu-S-MRC-5) cells by pre-pandemic plasma (Figure 1a). Consistent with previous data (11), plasma from participants recovered from SARS-CoV-2 infection mediated weak S-specific ADCC (mean 7.0% lysis) with Abs from only 13/31 individuals inducing > 10% lysis (Figure 1b). Responses were weaker amongst those who were vaccinated and not previously infected, with only one participant mediating ADCC > 10% after one vaccination (Figure 1b). A second vaccination increased responses with 10/40 individuals demonstrating killing above 10% specific lysis (Figure 1b). Nevertheless S-specific ADCC remained low overall, comparable overall to that of participants with infection-induced immunity (mean 7.7% lysis; Figure 1b). In contrast, hybrid immunity substantially enhanced ADCC (mean 31.3% lysis) with the absolute number of participants mediating > 10% lysis increasing to 29/31 after a single vaccination (Figure 1b). There was no further increase to ADCC within the hybrid group upon second vaccination, however, one additional participant (30/31) mediated > 10% ADCC (Figure 1b).

**Figure 1.**
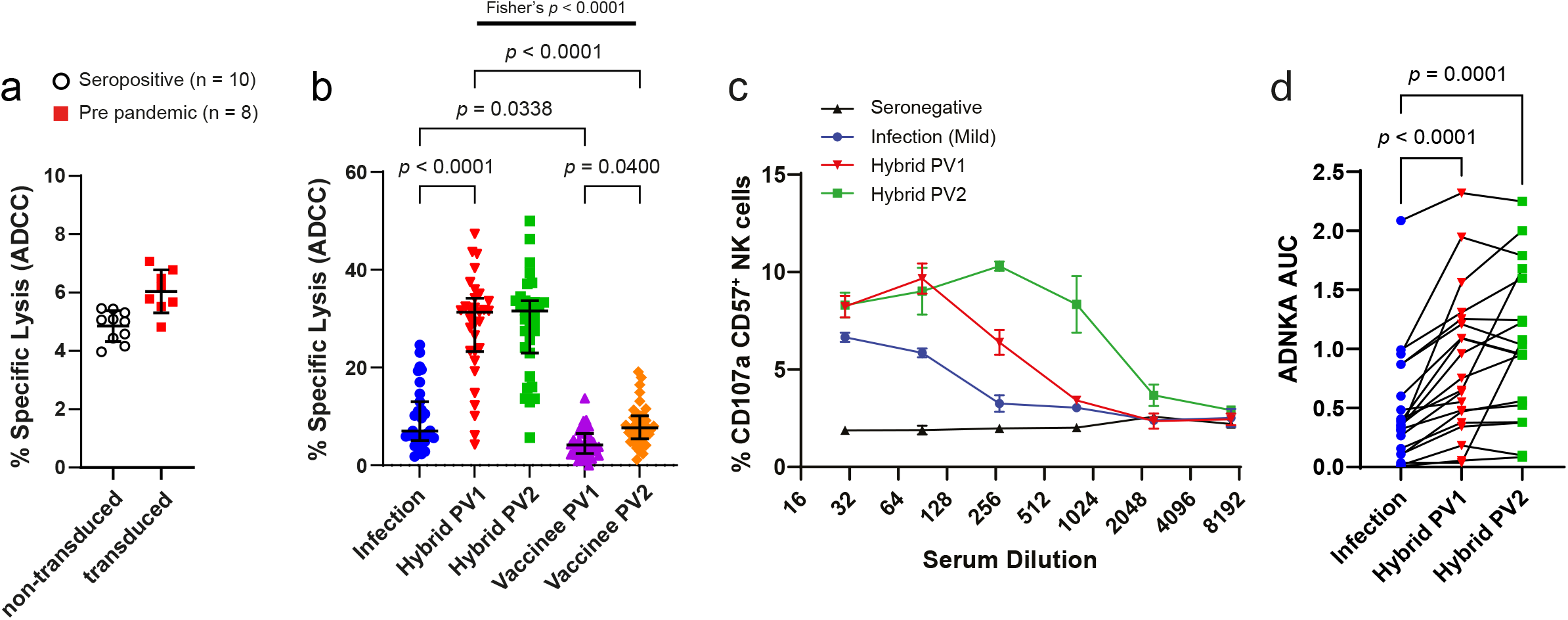
Vaccine-, infection-, and hybrid immunity-elicited S-specific NK cell activation. Background NK cell lysis of (a) non-transduced (open black circle) or Wuhan-Hu-1 S-expressing (red square) MRC-5 cells elicited by plasma from SARS-CoV-2 seropositive (n = 10) or plasma collected pre pandemic (n = 8), respectively, was measured by ^51^Cr release (E:T 25:1). (b) Sequential measures of S-MRC-5 cell ADCC elicited by plasma collected after infection then subsequent vaccination or after vaccination alone was measured by ^51^Cr release (E:T 25:1). Experiments were performed in duplicate with three independent donors and a representative plot shown. (c, d) Serum samples were serially diluted and CD57^+^ NK cell CD107a degranulation against A549-ACE2 cells infected with SARS-CoV-2 24 h previously at MOI = 5 was measured by flow cytometry. Data from a single individual is shown in (c) and compiled data from multiple donors in (d) was assessed by calculating the area under the curve (AUC). Black lines bisecting groups in (a) represent the mean of individual plasma samples with standard deviation, and in (b) represent median with IQR. *P* value in (b) was calculated using Kruskal-Wallis test with Dunn’s multiple comparisons test and in (d) calculated using one-way ANOVA for matched data with Tukey’s correction and shown above horizontal lines spanning comparison groups when significant. The probability of hybrid immunity inducing more robust ADCC than vaccine-induced immunity was calculated using two-sided Fisher’s exact test.

### Hybrid immunity induces significant S-specific ADCC against infected cells

Although cells transfected or transduced to express S provide a platform to isolate S-specific ADCC, Ab-dependent NK cell responses in the context of virus infection can differ significantly from that with cells overexpressing surface protein (11). As ^51^Cr release assays are incompatible with BSL3 conditions, we assessed Ab-dependent NK cell activation (ADNKA) against SARS-CoV-2 infected cells by measuring NK cell degranulation (CD107a) as a surrogate for ADCC.

During infection, ADNKA is dominated by Ab targeting antigens other than S, obscuring increases in S-specific ADNKA attributed to vaccination (11). To circumvent this issue, we focused on individuals with asymptomatic or mild infection (and, therefore, comparatively weak pre-vaccine ADNKA responses) followed by vaccination with a S subunit vaccine, and tested sera over a range of dilutions. This allowed us to select a dilution at which infection-induced anti-N/M/ORF3a responses had faded, but where a significant boost to S-specific ADNKA could be measured following vaccination (Figure 1c) across multiple donors (Figure 1d). Although a second vaccine dose enhanced the abundance of ADNKA-capable Ab in some donors, such that reactivity was maintained at higher dilutions (Figure 1c), it did not alter maximal levels of ADNKA (Figure 1c, d).

Therefore, anti-S Abs can mediate S-specific Ab-dependent NK cell activity against transduced or infected cells. However, these Abs are not efficiently induced by natural infection or vaccination – a hybrid combination is required.

### Neutralisation and ADNKA engage different Ab populations

We next used infected cells to compare S-specific ADNKA responses mediated by hybrid immunity induced by mild or severe infection to determine whether S-specific ADNKA provided by hybrid immunity approached the level of ADNKA induced by the sum of Ab targeting all SARS-CoV-2 cell-surface proteins *(i.e.* including anti-N/M/ORF3a; Figure 2a) (11). Individuals were further stratified by the vaccine they received [adenovirus (AstraZeneca ChAdOx1-S) or mRNA (Pfizer-BioNTech)]. Vaccination alone induced weak ADNKA, irrespective of vaccine platform (Figure 2a, b). In contrast, hybrid immunity boosted S-specific ADNKA to levels comparable to the potent multi-antigen ADNKA response of individuals who had recovered from mild infection, but not to the extent of those recovered from severe infection (Figure 2b). This pattern was in sharp contrast to the neutralising Ab response, where vaccination induced responses comparable to mild infection, and hybrid immunity gave responses comparable to severe infection (Figure 2c).

**Figure 2.**
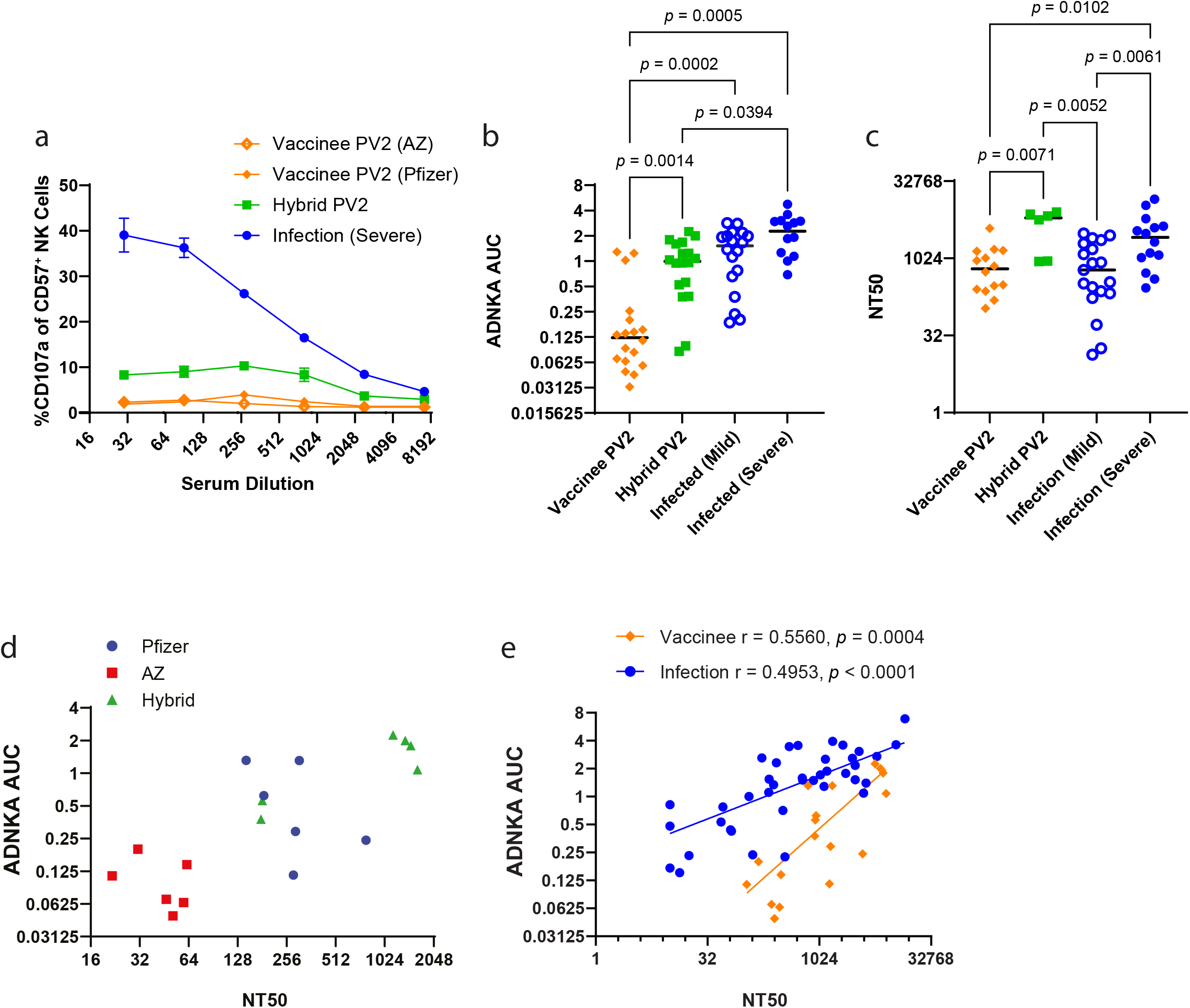
Spike-directed ADNKA against SARS-CoV-2-infected cells. A549-ACE2 cells were infected with SARS-CoV-2 for 24 h and CD57^+^ NK cell CD107a expression measured in response to serial dilutions of sera from vaccinees (PV2), hybrid immunity (PV2) or persons recovered from mild or severe infection and no vaccination. Representative data in (a) is depicted at the indicated dilutions and (b) the AUC was calculated, and data compiled for multiple donors (vaccinee n = 18; hybrid immunity n = 18; mild infection n = 18; severe infection n = 14). (c) Sera used in (b) were applied to SARS-CoV-2-infected cells, neutralisation assessed and NT50 calculated. (d, e) The significance of correlations between ADNKA and NT50 were assessed using Spearman’s correlation.

To investigate further, we considered the correlation between neutralising and ADNKA Ab titres against live virus. Comparisons amongst vaccinated individuals in which ADNKA was primarily driven by anti-S responses demonstrated a positive correlation between neutralisation and ADNKA, along with a clear hierarchy of responses. Vaccination with adenovirus (AstraZeneca ChAdOx1-S) demonstrated the weakest neutralisation and ADNKA activity, mRNA (Pfizer-BioNTech) vaccination induced better ADNKA and neutralisation than adenovirus vaccination, and both vaccination regimes were inferior to neutralisation and ADNKA induced by hybrid immunity (Figure 2d). Although neutralisation and ADNKA showed positive correlations in vaccinees and infected individuals (11), the relationship between the two activities was markedly different. For any given level of neutralisation, infection showed superior levels of ADNKA compared to vaccination (Figure 2e). Thus, in comparison to infection, S-based vaccines are effective at inducing neutralising Abs and less so at inducing ADNKA, consistent with the presence of Abs against multiple different ADCC targets following infection.

### Superior ADCC from hybrid immunity is not strictly explained by Ab abundance

To test whether differences in ADCC between hybrid and vaccine-induced immunity were due in part to IgG abundance, we compared ADCC with measurements of total IgG targeting full length S (FLS) and individual S1 and S2 subunits. Comparisons were also carried out on IgG3 subclass Ab; although IgG1 is most abundant, IgG3 supports more potent ADCC (36, 37).

ADCC positively correlated with amounts of anti-S1 and S2 IgG (Figure 3a) and IgG3 (Figure 3b) Abs, despite anti-S1 IgG3 not being abundant in those recovered from infection (Figure 3b). As the hybrid group demonstrated more robust ADCC after first vaccination and there were no significant increases in the levels of Abs or ADCC after second vaccination (see below), we focused on responses elicited from the hybrid group after one vaccine to contrast with the vaccinees post second vaccination. In both the hybrid cohort (PV1) and the vaccine group (PV2), ADCC correlated with increased anti-FLS IgG Ab levels (Figure 3c). This trend persisted when we compared anti-S1 IgG (Figure 3d) and anti-S2 IgG (Figure 3e) Ab levels with ADCC. There was also a significant correlation between levels of anti-FLS IgG3 Ab and ADCC in both the hybrid and vaccine cohorts (Figure 3f), although this correlation was lost for the hybrid group when comparing reactivity against individual S1 and S2 subunits of SARS-CoV-2 S (Figure 3g, h). This may reflect that the lower abundance of IgG3 results in IgG1 having a greater influence on total ADCC activity.

**Figure 3.**
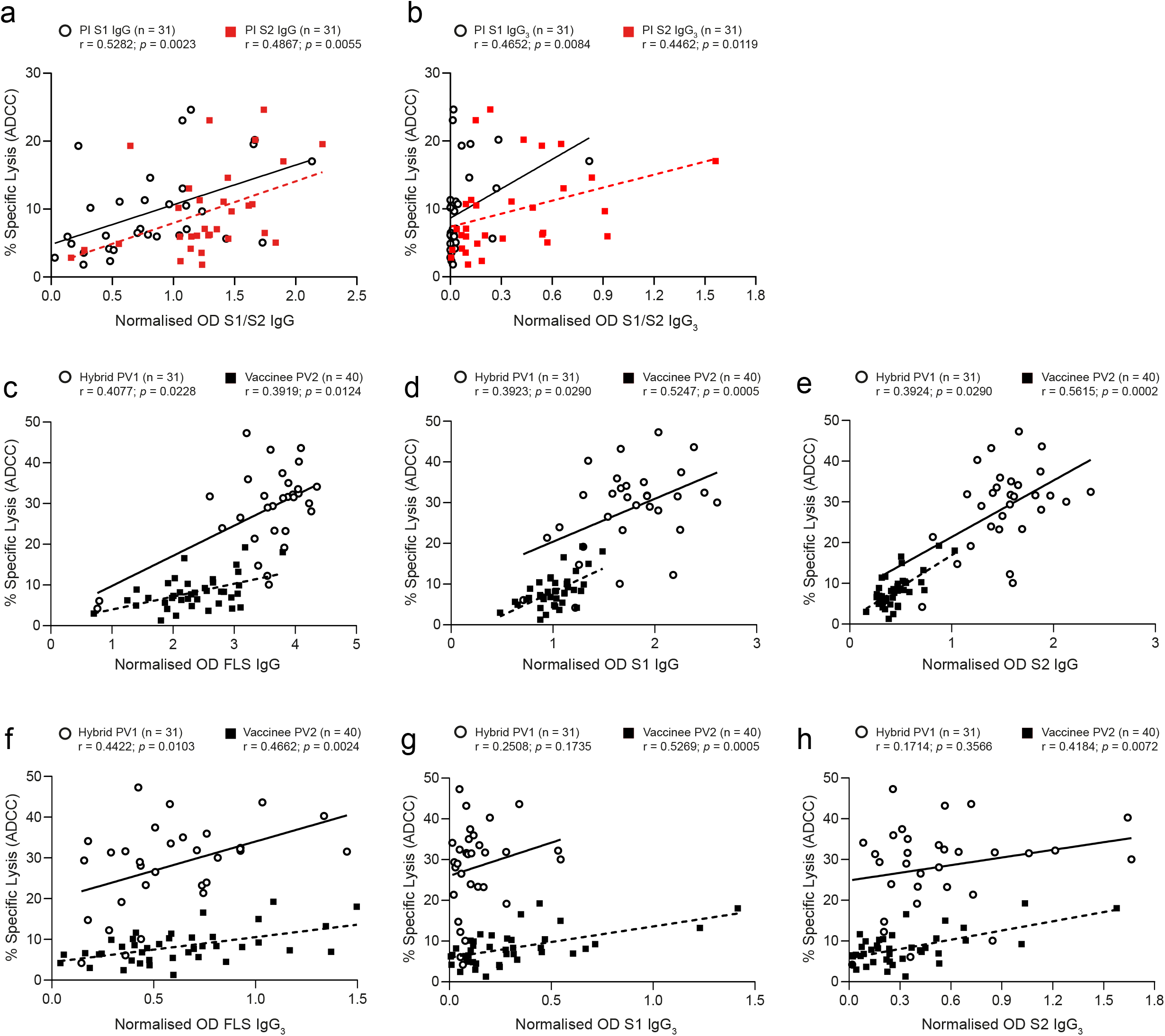
Associations between ADCC and anti-S1/S2 IgG and IgG3 Ab abundance. Correlations between ADCC induced by plasma Ab from samples collected post infection (PI) and levels of anti-S1 (open black circle) and anti-S2 (red square) (a) IgG and (b) IgG3 are plotted. Relationships between the magnitude of ADCC mediated by hybrid (PV1; open black circle) and vaccinee (PV2; black square) plasma Ab and (c) anti-FLS IgG, (d) anti-S1 IgG, (e) anti-S2 IgG or (f) anti-FLS IgG3, (g) anti-S1 IgG3 and (h) anti-S2 IgG3 are depicted.

Despite these strong correlations between Ab levels and ADCC *within* cohorts, there were significant differences *between* the cohorts, with Abs from vaccinees eliciting much weaker ADCC for a given level of Ab compared to their hybrid counterparts. Thus, although Ab levels influence ADCC activity, Ab abundance alone does not explain the superior ADCC induced by hybrid immunity.

### Vaccination, infection, and hybrid immunity induce distinct antibody responses to S1 and S2

Since strong ADCC induced by hybrid immunity reflects the quality of immune response rather than simply abundance of IgG, we investigated the specificity of anti-S Abs following vaccination or hybrid immunity in more detail. Circulating IgG Abs against FLS and the individual S1 and S2 domains were measured by ELISA after infection, and after first and second vaccinations. Anti-FLS IgG Abs were detected (OD > 0.1) from 30/31 of the participants included in the hybrid cohort, and 38/40 of the vaccinated individuals (Figure 4a). Within the hybrid cohort, levels of anti-FLS Abs rose significantly after first vaccination, but there was no further increase upon second vaccination (Figure 4a). In contrast, Ab levels for the vaccinee cohort increased significantly following the second vaccination but remained relatively low (Figure 4a).

**Figure 4.**
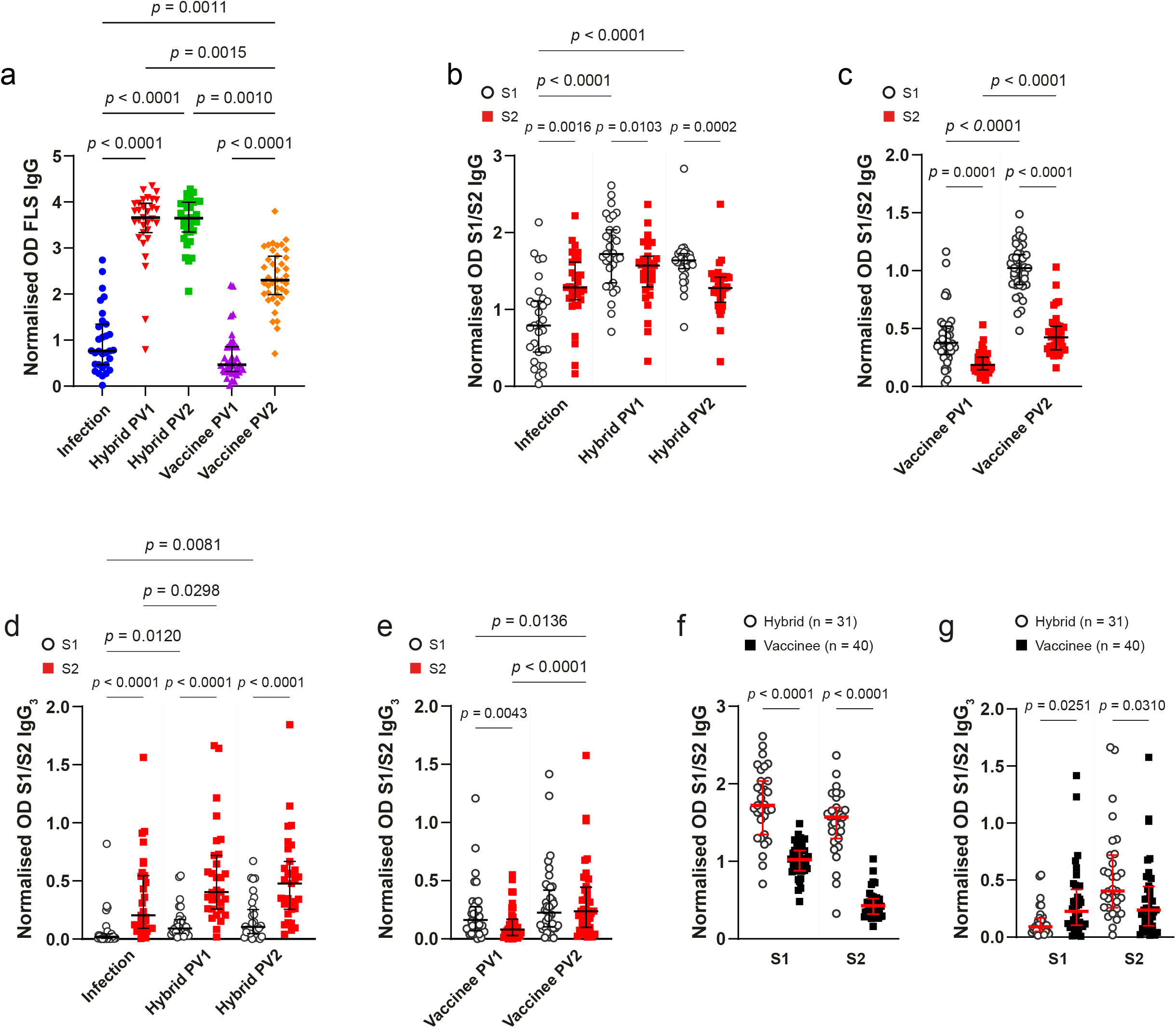
Anti-S1/S2 Ab responses after vaccination, infection, and hybrid immunity. Circulating anti-FLS IgG Abs from (a) participants with hybrid immunity were measured post infection (PI; blue circle) and after first (PV1; red chevron) and second (PV2; green square) vaccinations, and from vaccinated participants (n = 40) PV1 (purple triangle) and PV2 (orange diamond). Levels of anti-S1 IgG (open black circle) and anti-S2 IgG (red square) in samples obtained from (b) participants with hybrid immunity were compared PI, PV1 and PV2, and from (c) vaccinees were compared PV1 and PV2. Relative amounts of anti-S1 and anti-S2 IgG3 were measured for (d) persons with hybrid immunity or (e) vaccinees and anti-S1 and anti-S2 IgG3 Ab levels compared. Hybrid- and vaccine-induced anti-S1 and anti-S2 (f) IgG and (g) IgG3 were contrasted. P values in (a) were calculated using Kruskal-Wallis test with Dunn’s multiple comparisons test; (b) – (e) using Friedman’s test with Dunn’s multiple comparisons test; (f), (g) using Kruskal-Wallis test with Dunn’s multiple comparisons test and are shown above horizontal lines spanning comparison groups when significant. Lines bisecting groups represent median with IQR.

Infection generated IgG Abs against both S1 and S2 domains, however, anti-S2 Ab responses were favoured over anti-S1 (Figure 4b). Vaccination after infection increased anti-S1 IgG Ab levels but had no significant impact on the levels of anti-S2 IgG Abs, indicating preferential boosting of anti-S1 Ab (Figure 4b). Similar skewing towards anti-S1 responses following vaccination was also apparent amongst those vaccinated with no prior infection, as substantially higher levels of anti-S1 IgG Abs compared with anti-S2 IgG Abs arose after both first and second vaccinations (Figure 4c). Thus, the anti-S IgG Ab response differs following infection or vaccination, with vaccination selectively inducing anti-S1 IgG Abs, while infection-induced immunity promoted higher levels of anti-S2 IgG Abs.

### Infection history dictates anti-S2 IgG3 antibody responses

In addition to measuring the proportion of total IgG targeting S, we assessed the prevalence of IgG3 Abs in persons with hybrid or vaccine-induced immunity. Anti-S1 IgG3 Ab levels after infection were generally low, with only 4/31 participants having OD > 0.1 (Figure 4d), but levels increased after first vaccination with anti-S1 IgG3 Abs detected in 11/31 participants (Figure 4d). In contrast, anti-S2 IgG3 Ab levels were more substantial following infection, with 21/31 of participants having detectable anti-S2 IgG3 Abs (Figure 4d) and vaccination further improved anti-S2 IgG3 Ab levels (Figure 4d). With hybrid immunity, levels of anti-S2 IgG3 Abs were significantly higher than anti-S1 IgG3 Abs and remained stable up to ~5 months.

Vaccination alone resulted in 28/40 individuals producing anti-S1 IgG3 Abs after first vaccination and 31/40 after second vaccination (Figure 4e). One vaccination induced low amounts of anti-S2 IgG3 Abs in 16/40 persons that improved upon second vaccination with 30/40 participants having detectable anti-S2 IgG3 Abs (Figure 4e). Compared with hybrid immunity, levels of anti-S1 IgG3 Abs outweighed anti-S2 IgG3 following a single vaccination, with levels similar following two vaccinations (Figure 4e). Therefore, for both total IgG and IgG3, infection-induced immunity favoured anti-S2 Ab production, whereas vaccination induced more robust anti-S1 Ab levels (Figure 4f, g). The strongest IgG Ab responses against both S1 and S2 were seen following hybrid immunity.

### ADCC potency depends on antibodies reactive against both S1 and S2

To determine whether Abs targeting either or both of the S1 and S2 domains were responsible for potent S-specific ADCC following hybrid immunity, we selectively depleted Abs that target different S domains and assessed ADNKA against infected cells. ADNKA induced by vaccinee serum was too low to observe clear effects following depletion so we focussed solely on donors with hybrid immunity. Using two donors, we depleted Abs targeting the entire S trimer, the S1 domain, the S2 domain, or specific NTD or RBD domains found within S1. Serums were pre-diluted such that the starting dilution was sufficient to have diluted out Abs targeting antigens other than S (Figure 1c). Although depleting Abs targeting NTD or RBD alone led to only minor reductions in ADNKA, complete S1 or S2 subunit Ab depletion led to significant reductions in both donors (Figure 5a, b), indicating that a combination of Abs targeting S1 and S2 is required for robust ADNKA.

**Figure 5.**
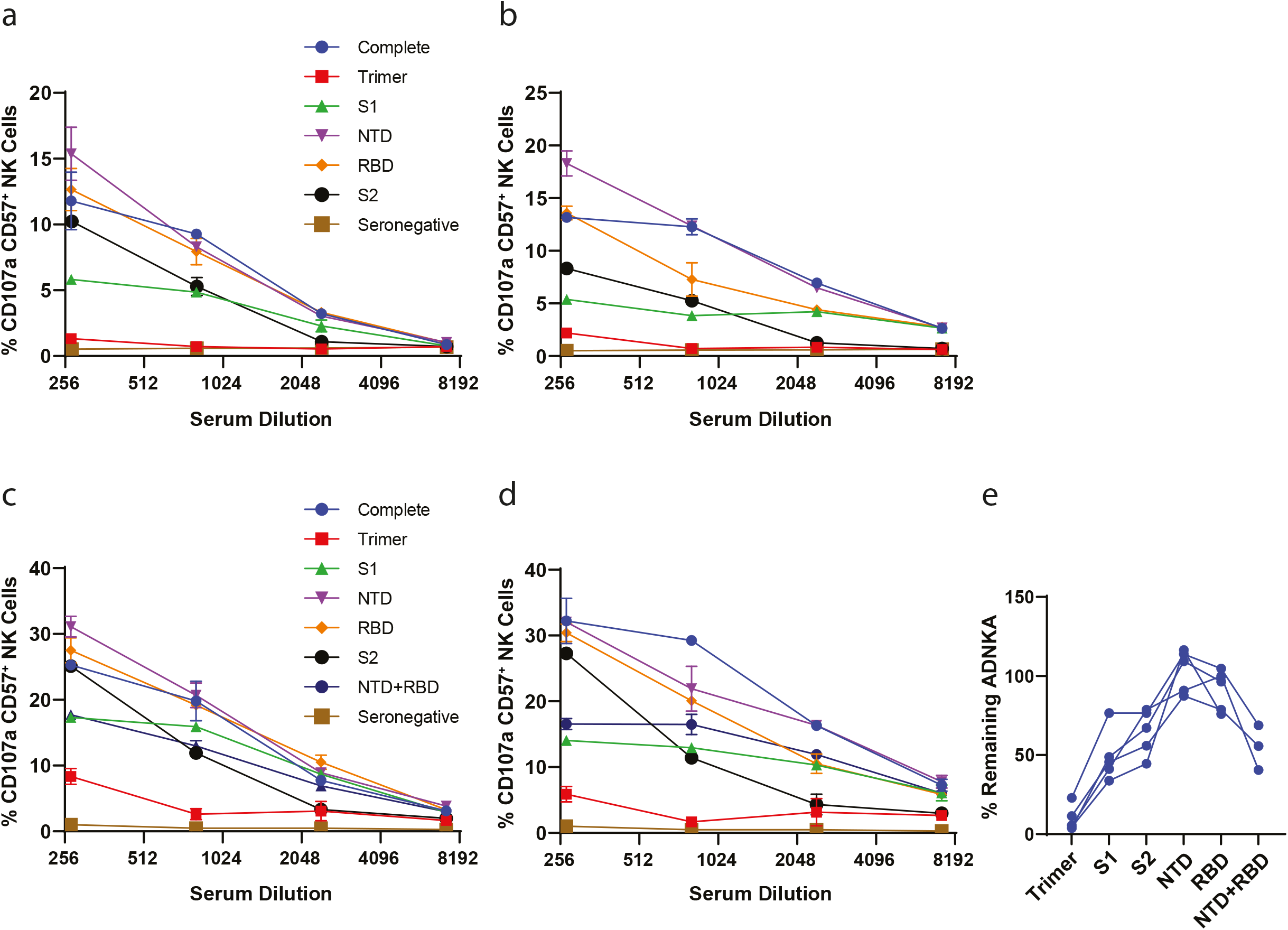
Assessment of ADNKA after S subunit Ab depletion. Sera from individuals with hybrid immunity were diluted 1:9 then Abs targeting the indicated domains of S were depleted using magnetic bead-conjugated protein. These depleted serums were then used to measure CD57^+^ NK cell CD107a expression at the indicated dilutions in the presence of SARS-CoV-2-infected A549-ACE2. Four individuals are depicted in (a) – (d). (e) The AUC for each sample in (a) – (d) was calculated then the amount of ADNKA remaining following depletion of Abs targeting each domain in relation to sera containing all Abs, or sera from which all S Abs had been depleted, was determined (n = 5).

Depleting Abs targeting NTD or RBD yielded smaller reductions in ADNKA than depleting Abs targeting the entire S1 domain. The reagents used for NTD and RBD depletion targeted amino acids 16-318 and 319-541, respectively, while the S1 depletion targeted amino acids 16-685. As the S1 depletion potentially removed Abs that targeted 542-685, which were not depleted by RBD or NTD depletion, we repeated this experiment and included a double depletion of both NTD and RBD (Figure 5c, d). The RBD/NTD double depletion closely mirrored S1 depletion. Combined data from five donors indicated that potent ADNKA induced by hybrid immunity was dependent on the presence of Abs reactive to both the S1 and S2 regions, as depletion of either region led to a significant loss in ADNKA (Figure 5e).

### Antibody reactivity against three determinants along S correlates with potent ADCC

Having determined that Abs against both S1 and S2 subunits were important for eliciting strong S-specific ADCC, we investigated the fine specificity of Ab responses arising from hybrid immunity by ELISA-based peptide scanning. 181 overlapping peptides were coated onto ELISA plates in sequential pairs and IgG Ab reactivity was measured. A comparison of Abs reactive to linear regions contained in both S1 and S2 produced after infection (Figure 6a, blue line) and subsequent vaccination (Figure 6a, orange line) revealed differential patterns and robustness of Ab reactivity. A heatmap illustrating changing patterns of linear Ab epitope reactivity along S before and after vaccination is depicted for 7 participants in the hybrid group (Figure 6b).

**Figure 6.**
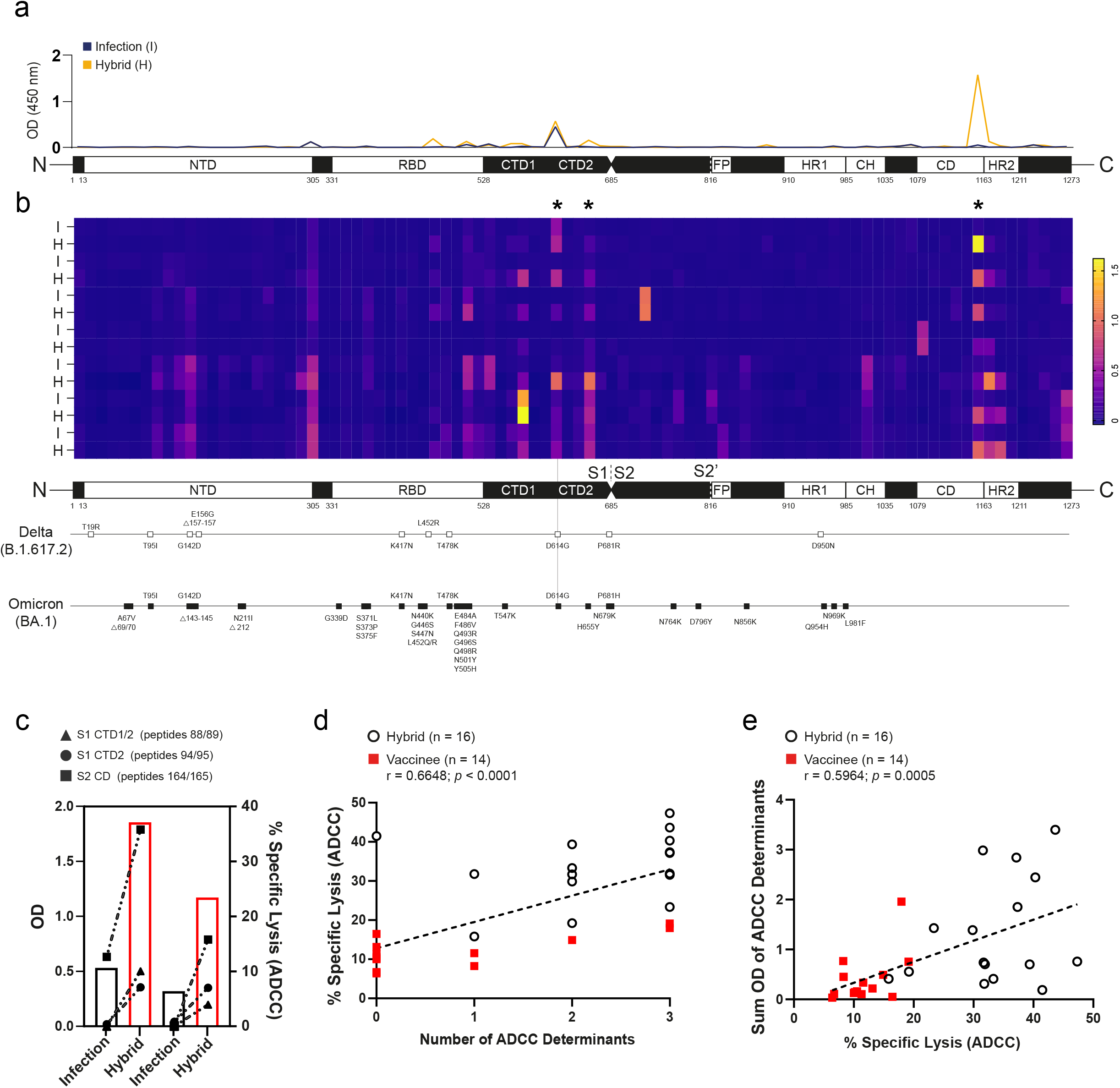
Peptide scanning to identify distinct linear regions associated with robust ADCC. Linear FLS Ab epitope reactivity was determined by ELISA-based peptide scanning. (a) A representative depiction of anti-IgG Ab reactive to linear segments is illustrated by an overlaid line graph. The blue line represents OD results of the full S peptide scan of a sample collected from one participant recovered from moderate COVID-19 infection and the orange overlay represents a sample from the same participant collected 1 month after their first vaccination. (b) Compiled peptide scan data from samples collected from 7 participants post infection (PI) and after hybrid immunity (H) were illustrated using a heat map and aligned with known mutations in Delta and Omicron sequences. Three determinants associated with ADCC are identified by asterisks. (c) The left axis depicts anti-S1 CDT1/2 (triangle), S1 CTD2 (circle) and S2 CD (square) IgG Ab OD and compares these levels with ADCC (right axis) for two participants after infection (open black bar) and after subsequent vaccination (open red bar). Participant IgG Ab OD collected from peptide scanning were (d) scored for number of distinct regions or (e) tallied and associations between Ab reactivity and levels of ADCC assessed.

Samples obtained after infection alone had Abs recognising fewer S1 and S2 epitopes than matched samples following vaccination (Figure 6b), with Ab reactivity against three distinct regions along S particularly enriched amongst the hybrid samples (Figure 6b). These three areas of reactivity are contained within the (i) C-terminal domain (CTD) 1 and (ii) CTD2 of S1 and (iii) a region in S2 immediately upstream of the heptad repeat 2 (HR2) sequence in the connector domain (CD; Figure 6b). Besides the early arising and common D614G mutation, these regions showed no genetic variation between Delta or Omicron variants (Figure 6b).

To illustrate potential associations between ADCC and Ab reactivity against these three regions, we used data collected from two persons with infection-induced immunity and subsequent vaccination to depict the relationship between ADCC and the magnitude of anti-IgG Ab found in CTD1/2 and CD. In both cases, increasing anti-IgG Ab levels specific for each of the three regions paralleled robust increases in ADCC (Figure 6c). Further, there were significant associations between ADCC and both the number of ADCC determinants to which a participant demonstrated IgG Ab reactivity (Figure 6d) and the cumulative OD from the three distinct regions (Figure 6e). Thus, epitope specificity plays a key role in S-specific killing of SARS-CoV-2-infected cells.

### Hybrid immunity broadly mediates ADCC across variant strains

Virus evolution in human populations results in selection of viral variants mutated at key residues for neutralising Ab binding – primarily within the RBD, but also within the NTD (7, 8). However, ADCC-susceptible determinants are distributed much more broadly across S (Figure 5) and ADCC determinants revealed by peptide scanning do not overlap with mutations in recent virus variants such as Delta and Omicron (Figure 6b). This suggests ADCC induced by hybrid immunity may be better preserved than neutralisation across variant strains. To assess S-specific Ab-dependent responses against variant strains, MRC-5 cells were transduced to express Delta or Omicron S at levels equivalent to Wu-S-MRC-5 (Figure 7a). When used in ADCC assays, Abs produced after hybrid infection elicited comparable levels of ADCC against target cells expressing Wuhan-Hu-1 and Delta S (Figure 7b). Although ADCC was reduced against Omicron S-expressing cells, the relative reduction was mild (12.9% decrease), and 5/16 participants mediated equivalent levels of ADCC against cells expressing Wuhan-Hu-1, Delta and Omicron S (Figure 7b).

**Figure 7.**
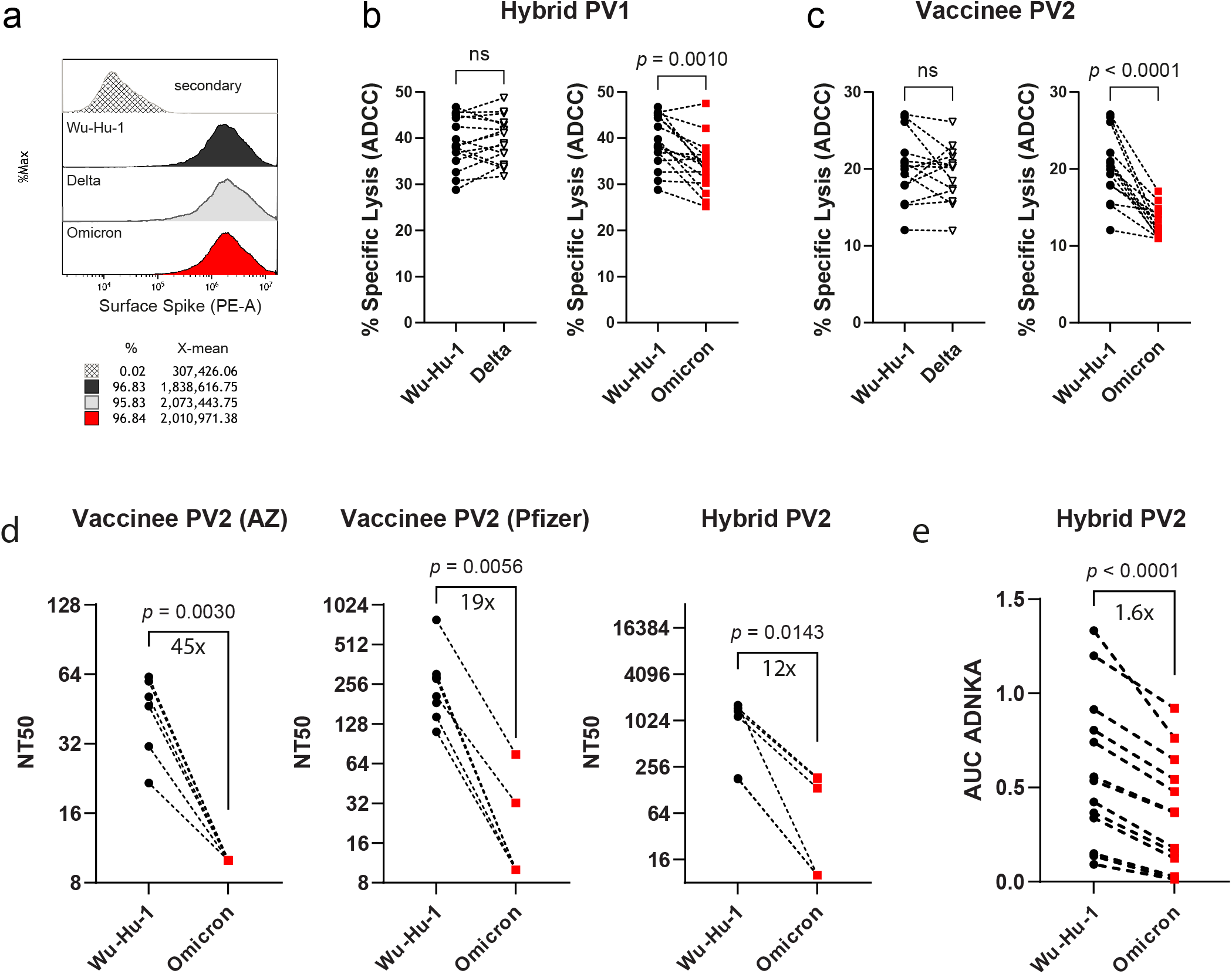
Vaccine- and hybrid-induced S-specific ADCC against variants of concern. (a) Histogram overlay of MRC-5 cells transduced to express similar levels of Wuhan-Hu-1 (black), Delta (grey) and Omicron (red) S protein used in ^51^Cr assay to assess the efficacy of Ab produced with (b) hybrid (n = 14) or (c) vaccine-induced (n = 16) immunity in eliciting ADCC against variant strains. (d) SARS-CoV-2 neutralisation assay was performed on serial dilutions of serum samples and NT50 values against either Wuhan-Hu-1 or Omicron-infected A549-ACE2 were calculated. (e) Serum samples were serially diluted and CD57^+^ NK cell CD107a expression against A549-ACE2 cells infected with either Wuhan-Hu-1 or Omicron was measured by flow cytometry and AUC calculated. P values in (b) - (e) were calculated using Student’s paired *t* test and shown above horizontal lines spanning comparison groups when significant.

A subset of vaccinees mediating detectable ADCC against Wu-S-MRC-5 were similarly tested. Here, the decline in ADCC against cells expressing Omicron S was more pronounced with a 35.5% relative reduction (Figure 7c). Ab from only 1/14 vaccinees mediated comparable levels of ADCC against wild type and variant strains (Figure 7c), consistent with Ab in these donors being focussed on the S1 domain, which is more heavily mutated in these virus variants.

To ensure that these results were comparable to an infection setting, we assessed neutralisation and ADCC against replicating virus using either ancestral Wuhan-Hu-1 or Omicron strains. Consistent with published data, we observed that Omicron was more resistant to neutralisation following vaccination, and despite this decrease in neutralisation being less severe following hybrid immunity compared with vaccination, a significant loss was observed (Figure 7d). In contrast, and in agreement with data obtained using cells overexpressing S, we measured only a small decline in ADNKA against Omicron-infected cells (Figure 7e). Thus, by targeting a broader range of epitopes, including those more conserved across variants, ADCC resists virus escape due to virus mutation more effectively than neutralisation.

## Discussion

Strong and durable immune responses are desirable to limit SARS-CoV-2 transmission and control severity of infection. Neutralising Ab activity has dominated as a surrogate measure of protection, however, protection against severe illness without robust neutralisation suggests that other aspects of immunity, including T cells and NK cells, play a role (38). The ability of T cells and NK cells to limit illness by eliminating infected host cells once infection does occur underlies the importance of considering their recruitment in vaccination strategies, especially if their activity is better conserved across variants than Ab neutralisation (39, 40).

Associations between ADCC and viral control in animal models (12–22) indicate that it is an important component of immunological protection. Infection with SARS-CoV-2 induces Abs against N/M/ORF3a that dominate NK cell activation by infected cells (11), however, in the absence of vaccines encoding alternative ADCC antigens, it is important to understand the capacity for S-specific ADCC. Infection or vaccination alone was insufficient to induce potent S-directed ADCC, whether assessed by direct killing of S-expressing cells or NK cell activation against SARS-CoV-2 infected cells. However, the strong ADCC observed with hybrid immunity indicates that it is possible to generate robust and lasting ADCC through S-targeted Ab responses.

Previous analysis of hybrid immunity in the context of neutralising Abs demonstrated that compared to vaccination alone, it results in more memory B-cells and circulating Abs, the latter of which was affirmed in our study (41, 42). This increase in circulating IgG enhances neutralisation of more immune-evasive virus variants (41, 43, 44), while at the clonal level, it also reflects selection of Abs with higher neutralising potency (45). In the case of ADCC, neither increased abundance, nor differences in IgG subclass could explain superior hybrid immunity-induced ADCC. Instead, qualitative aspects of the Ab profile played a more significant role. Robust levels of Abs targeting numerous epitopes in both S1 and S2 were a consistent feature of hybrid immunity.

The S1 domain of spike contains the NTD and RBD domains, which are required for receptor binding and are dominant targets for neutralising Abs. As a result, vaccine development has focussed heavily on inducing S1-targeted Abs, including the testing of RBD subunit vaccines (46–50). However, due to mutations selected in S1 that tend to reduce neutralisation, there is increasing interest in neutralising Abs targeting the more conserved S2 domain (51). Studies in mice demonstrated that S2-specific vaccination effectively induces cross-variant neutralisation (51). Our work now extends this by showing that strategies to induce both S1 and S2 responses will also serve to boost ADCC. The polyspecific *(i.e.* S1 and S2) nature of S-directed ADCC improves targeting of virus variants with heavily mutated S1 and may limit the ability of viruses to evade this arm of host defense. With a wider range of Abs recognizing epitopes distributed throughout S, the loss of any single epitope may not substantially impact ADCC, raising the barrier for selection of escape mutants.

The S gene of seasonal human beta coronaviruses (HCoV HKU1, HCoV OC43) is divergent from SARS-CoV-2, sharing only ~ 36% overall homology, with S2 somewhat more conserved than the S1 domain. The preferential Ab targeting of S2 epitopes after infection may be partly attributed to a heterologous boost towards cross-reactive S2-specific memory B cells (52–54), with pre-existing cross-reactive HCoV memory B cells activated during SARS-CoV-2 infection kick-starting production of anti-S2 Abs. Interestingly, the same regions we revealed as key regions of Ab reactivity corresponding with potent ADCC were previously demonstrated to be elicited by heterologous anti-S Ab responses (54). This raises the question as to whether immunodominant regions within S1 and S2 conserved between HCoV and SARS-CoV-2 are selectively targeted for Ab reactivity and avidity to support robust ADCC. If SARS-CoV-2 infection induces cross-reactive HCoV anti-S2 Ab production, it is unclear why vaccination did not have a similar effect.

Prolonged antigen exposure from high viral loads, the presence of inflammatory stimuli during virus infection, the way S traffics to and is presented on the infected cell surface, or differential sites of antigen presentation (e.g. intramuscular for the vaccine, respiratory for the virus), may all contribute to diverse outcomes from exposure through infection versus S subunit vaccination. It will be interesting whether switching vaccine delivery from intra-muscular to intranasal, to promote mucosal immunity, impacts this process (55, 56).

This study focussed on individuals who were infected prior to vaccination. Many individuals have now been infected after being vaccinated with ancestral S antigen and it is unclear how these breakthrough infections will affect ADCC against emerging variants. Our previous research demonstrated that infection induces non-S ADCC responses that target N/M/ORF3a (11). However, as vaccination reduces the induction of non-S (e.g. nucleocapsid) Ab during breakthrough infection, presumably due to enhanced vaccine-mediated immunological control (57–59), breakthrough infections may fail to induce potent N/M/ORF3a-mediated ADCC. In this case, strong S-directed ADCC induced by hybrid immunity would become an important effector mechanism contributing to protection and an important consideration for future research.

In summary, our data reveal that hybrid immunity establishes conditions whereby Abs generated against key determinants within S1 and S2 domains elicit ADCC quantitatively superior to vaccination or infection alone. In addition, hybrid immunity induced by ancestral antigen engenders an Ab response that retains activity against variant strains to a far greater extent than neutralisation. Given the complementary roles of neutralisation and ADCC in controlling cell-free and cell-associated virus, respectively, both Fc- and Fab V-region mediated effector functions are desirable for incorporation into vaccine strategies. Strong ADCC may play a part in the protection offered by hybrid immunity, uncovers a novel role for S2-targeted Abs, and suggests that vaccine strategies based on spike expression would benefit from inducing Abs targeted broadly across spike, as opposed to just RBD.

## Methods

### Study subjects

This study was carried out at two sites. In Canada, the study conformed to recommendations of the Canadian Tri-Council Policy Statement: Ethical Conduct for Research Involving Humans, and ethical approval was given by the Health Research Ethics Authority of Newfoundland and Labrador (HREB). Peripheral blood was collected from study subjects at approximately three-month intervals and a questionnaire addressing previous testing history and reasons for suspecting infection with SARS-CoV-2 was administered at study intake after written informed consent in accordance with the Declaration of Helsinki. Thirty-one individuals with confirmed infection who continued in the study and received 2 doses of a COVID-19 vaccine were matched with 40 individuals with no previous infection history who had received 2 doses of a COVID 19-vaccine (Table 1). Asymptomatic participants were identified through Public Health surveillance and contact tracing after contact with RT-PCR-confirmed cases or through serological testing for anti-S and anti-N protein IgG Abs (60). Most participants enrolled following the first wave of COVID-19 and infections were attributed to ancestral Wuhan-Hu-1 SARS-CoV-2. Three participants had infections attributed to B.1.1.7. Participants self-declared any medical treatments they were receiving as well as information on comorbidity. Persons with any known underlying immune compromising condition or on immunosuppressive treatment were excluded.

**Table 1.**
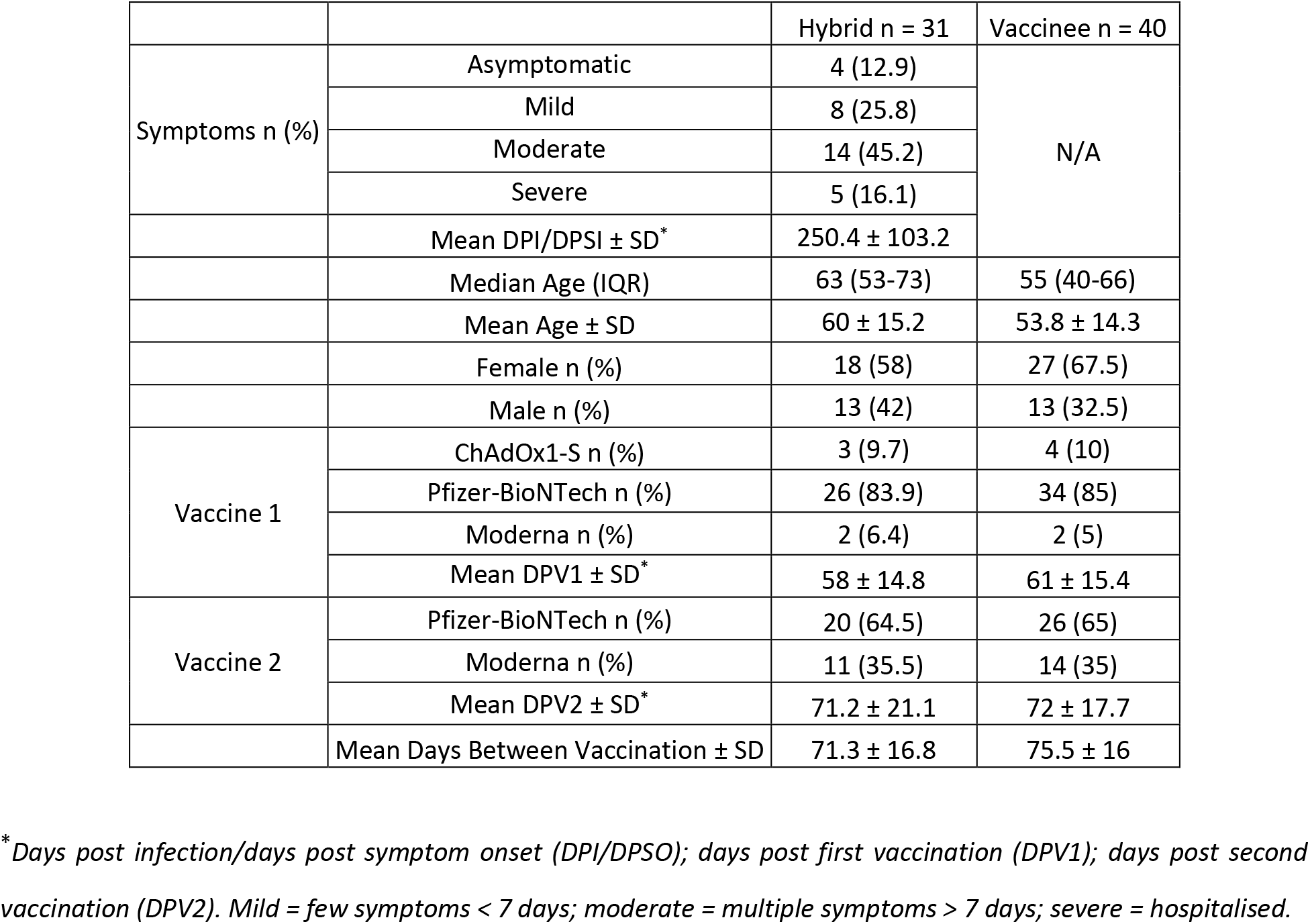
Relevant features and categorization of study cohort.

In the UK, ethical approval was given by Local Research Ethics Committee or the Avon Longitudinal Study of Parents and Children (ALSPAC) Ethics and Law Committee. Peripheral blood was collected 2-4 weeks after a RT-PCR or LFD confirmed infection, or after vaccination, from otherwise healthy donors. Alternatively, serum samples submitted as part of the ALSPAC cohort (61–63) were assessed based on serology for N and S, to identify individuals who had submitted samples after an initial infection, followed by samples after each vaccination. In total 18 individuals who were infected prior to vaccination were matched with 18 individuals who had received two doses of vaccines with no experience of prior infection. Results were also compared to samples from a previously described cohort of individuals who had experienced mild or severe COVID-19 and had not been vaccinated (11).

### Blood sample processing

Whole blood was collected by venipuncture in acid citrate dextrose vacutainers, after which plasma was collected following 10 min centrifugation at 500*g* and stored at −80°C (Canada). Alternatively, whole blood was collected in a serum-separating vacutainer and serum collected following centrifugation at 500*g* for 10 min (UK). PBMC used for ADCC experiments were isolated from anti-coagulated heparinized blood from healthy donors by density gradient centrifugation using the Canadian Autoimmunity Standardization Core consensus standard operating procedure (version: March 21, 2019). Freshly isolated PBMC were resuspended in lymphocyte medium consisting of RPMI-1640 with 10% FBS (HyClone™), 200 IU/mL penicillin/streptomycin, 0.01 M HEPES, 1% L-glutamine (all from Invitrogen) and 2 x 10^-5^ M 2-mercaptoethanol (Sigma-Aldrich) and used directly in functional experiments.

### Cell lines and viruses

All cell lines and PBMC were cultured with 5% CO_2_ at 37°C. VeroE6 cells expressing ACE2 and TMPRSS2 (VAT) and A549 cells expressing ACE2 (AA) were a gift from the University of Glasgow Centre for Virus Research (64). Human lung fibroblast MRC-5 cells were obtained from ATCC® (CCL-171™) and Lenti-X™ 293T cells were obtained from TakaRa (Mountain View CA, USA). All were propagated in complete DMEM (Sigma-Aldrich) containing 10% FBS (HyClone™) and 200 IU/mL penicillin/streptomycin (Invitrogen, Carlsbad CA, USA). Ancestral SARS-CoV-2 was recovered from a BAC containing the complete genome of a strain that matches the original Wuhan-Hu-1 (64) and the Omicron (BA.1) variant was a gift from Professor Arvind Patel (from the University of Glasgow Centre for Virus Research). Both were propagated in VAT cells. Virions were concentrated and purified by pelleting through a 30% sucrose cushion and titrated by plaque assay in AA cells as previously described (11). All virus seed stocks were verified by whole-genome sequencing on the Illumina platform.

### Spike gene transfer and expression in MRC-5 cells

The recombinant Lenti-X™ pLVX-IRES lentiviral vector expression system (TakaRa) was used to introduce Wuhan-Hu-1, Delta (B.1.617.2), or Omicron (BA.1) S sequences into the MRC-5 cell line. The Wuhan-Hu-1 S was obtained from BEI Resources in a pcDNA™3.1(-) mammalian expression vector (NR-52420; NIAID, NIH) (65). Delta S was synthesised by Invitrogen GeneArt (Thermo Fisher Scientific) and contains mutations: T19R, T95I, G142D, E156G, E157- F158- R450L, K476T, G612D, R679P, N948D. Omicron S was obtained from BEI Resources in a pCMV/R mammalian expression vector (NR-56470; NIAID, NIH) with the following mutations: A67V, H69-, V70-, T95I, G142-, V143-, Y144-, Y145D, N211-, L212I, “EPE” insertion between 214R and 215D, G339D, S371L, S373P, S375F, K417N, N440K, G446S, S477N, T478K, E484A, Q493R, G496S, Q498R, N501Y, Y505H, T547K, D614G, H655Y, N679K, P681H, N764K, D796Y, N856K, Q954H, N969K, L981F.

Constructs were inserted into pLVX-IRES as detailed in (66) using conventional cloning protocols and all constructs were verified by forward and reverse strand sequencing (TCAG Facilities, Hospital for Sick Children, Toronto ON, Canada) to ensure authenticity. SnapGene Software (San Diego CA, USA) was used for designing and visualizing cloning procedures, designing and aligning sequencing primers and comparing variant sequences to wild-type Wuhan-Hu-1. Transduced cells were propagated and selected in complete DMEM containing 1 μg/mL puromycin (Sigma-Aldrich). Extracellular S expression was confirmed by flow cytometry, as previously outlined (66).

### Antibody-dependent cell-mediated killing assays

PBMC were freshly processed, resuspended in lymphocyte medium and kept at 37°C and 5% CO_2_ until use. For ADCC assays using Wuhan-Hu-1, Delta or Omicron S-expressing MRC-5 target cells, 10^4^ cells/well were plated and labelled with 1 μCi Na_2_^51^CrO_4_/well (PerkinElmer, Akron, OH, USA) overnight in 96-well round bottom plates and then washed four times in PBS containing 1% FBS (HyClone™). PBMC (E:T 25:1) and heat-inactivated plasma (56°C for 1h) were added to wells with a final volume of 300 μL and final plasma dilution of 1:1000 and cytotoxic activity measured by ^51^Cr release over 5 h. ^51^Cr release was measured in 125 μL of supernatant on a Wallac 1480 Wizard gamma counter and percent specific lysis calculated by (experimental ^51^Cr release – spontaneous ^51^Cr release) / (maximum ^51^Cr release – spontaneous ^51^Cr release) x 100.

Assays using virus-infected cells used CD107a degranulation ADNKA as a proxy for ADCC to be in compliance with BLS3 containment and were carried out as previously described (11). Briefly, AA cells were infected at MOI = 5 for 24 h prior to being detached with TrypLE™ (Thermo Fisher, Waltham, MA USA), then 25,000 targets mixed with 250,000 PBMC, serum, and anti-CD107a-FITC (BioLegend, London, UK) and GolgiStop (BD, Swindon, UK) in a total volume of 100 μL for 5 h. PBMC were stained with live/dead fixable aqua (Thermo Fisher), anti-CD3-PE-Cy7 (UCHT1, BioLegend), anti-CD56-BV605 (5.1H11, BioLegend), and anti-CD57-APC (HNK-1, BioLegend). Data were acquired and analysed using an Attune NXT Flow Cytometer (Thermo Fisher) and expressed as the percentage of live CD107a^+^ CD57^+^ NK (CD3^-^ CD56^+^) cells. All sera were tested against mock-infected cells to ensure there was no background NK cell activation and any sera demonstrating background activation were excluded. A seronegative serum was included in all assays as a negative control. To enable comparisons with previous datasets, and to minimise inter-experiment variability, a donor serum demonstrating moderate ADNKA was included as a positive control in every assay. Sera were tested at a range of dilutions, and the area under the curve (AUC) was calculated using GraphPad Prism 9. This value was then normalised to the AUC for the standard serum in each assay.

### Virus neutralisation assay

Assays were carried out as previously described (11). Briefly, 600 plaque forming units of SARS-CoV-2 was incubated with appropriate dilutions of serum, in duplicate, for 1 h at 37°C. The mixes were then added to pre-plated VeroE6 cells for 48 h. After this time, monolayers were fixed with 4% paraformaldehyde (Thermo Fisher), permeabilised for 15 min with 0.5% NP-40 (Merck Life Science, Dorset, UK) then blocked for 1 h in PBS containing 0.1% Tween (PBST) and 3% non-fat milk. Primary Ab (anti-N 1C7, Stratech, 1:500 dilution) was added in PBST containing 1% non-fat milk and incubated for 1 h at room temperature. After washing in PBST, secondary Ab (anti-mouse IgG-HRP, Pierce, 1:3000 dilution) was added in PBST containing 1% non-fat milk and incubated for 1 h. Monolayers were washed again, developed using Sigmafast™ OPD (Sigma-Aldrich) according to manufacturers’ instructions, and read on a Clariostar Omega plate reader (OD 450 nm). Wells containing no virus, virus but no Ab, and a standardized serum displaying moderate activity were included as controls in every experiment. NT50 were calculated in GraphPad Prism 9.

### Serological testing

Plasma was diluted in PBS containing 0.05% TWEEN® 20 (0.05% PBST; Sigma-Aldrich, St. Louis, MO, USA) and 0.1% bovine serum albumin (BSA, Sigma-Aldrich) then tested against recombinant proteins coated in Dulbecco’s PBS (DPBS, Sigma-Aldrich) overnight onto 96-well Immunlon-2 plates (VWR Scientific, Mississauga, ON, Canada). Recombinant protein antigens included SARS-CoV-2 FLS glycoprotein trimer [50 ng/well; SMT1-1 reference material, National Research Council (NRC), Canada], the S1 subunit of SARS-CoV-2 S (65 ng/well; SinoBiological, Wayne, PA, USA) and S2 subunit of SARS-CoV-2 S (50 ng/well; SinoBiological). The predicted molecular mass for S1 and S2 were 76.5 kDa and 59.4 kDa, respectively, and coating amounts were determined to account for this difference. Plates were washed 4 times with 0.05% PBST, blocked for 1 h with 200 μL of PBS containing 1% BSA (Sigma-Aldrich), washed 4 times then 100 μL/well of diluted plasma (1:500 for FLS IgG or 1:100 for S1 and S2 IgG and IgG3) was applied to antigen-coated plates in duplicate wells for 1.5 h. Total IgG was measured following 6 washes and a 1-h incubation with 100 μL/well of 1:50,000 HRP-conjugated polyclonal goat anti-human IgG (Jackson ImmunoResearch Labs, West Grove, PA, USA). IgG3 was measured following 6 washes and a 1-h incubation with 100 μL/well 1:5,000 mouse anti-human biotin-conjugated IgG3 hinge (SouthernBiotech, Birmingham, Alabama, USA) then 6 washes and 1-h incubation with 100 μL/well 1:40,000 HRP-conjugated streptavidin (Jackson ImmunoResearch Labs). Plates were developed using tetramethylbenzidine (TMB) substrate (Sigma Aldrich) following 6 washes then incubated in the dark at room temperature for 20 min. Reactions were stopped with an equal volume of 1 M H_2_SO_4_ and optical density (OD) read on a BioTek synergy HT plate reader at 450 nm.

### Antibody depletions

Depletions of specific Abs from sera were carried out as previously described (11). In brief, Abs targeting different domains of S were depleted using magnetic bead conjugated proteins based on either the RBD, NTD, entire S1, or entire S2 domain of SARS-CoV-2 Wuhan-Hu-1 S (ACROBiosystems). Beads were resuspended in PBS + 0.05% BSA, then serum was pre-diluted 1:9, and 50 μl mixed with 150 μl beads. Mixtures were incubated on a rotating mixer at 4 °C overnight. Serum diluted in buffer alone was used as a control. Magnetic beads were then removed using a 3D printed magnetic stand then a second round of depletion carried out using fresh beads as indicated. All values in assays were corrected for dilutions.

### Peptide scan ELISA

Individual overlapping peptides (17- or 13-mers, with 10 amino acid overlaps) spanning the canonical Wuhan-Hu-1 S sequence (NR-52402; BEI Resources) were reconstituted at 10 mg/mL in DMSO (Sigma-Aldrich) then diluted to 50 μg/mL in DPBS (Sigma-Aldrich) and stored at −20°C. 125 ng/well of each S peptide 2 – 181 (BEI Resources) was coated overnight on Immunlon-2 plates (VWR Scientific) in sequential pairs (e.g. 2 & 3… 180 & 181). Full length S trimer (SMT1-1, NRC) was diluted in DPBS and coated overnight at 150 ng/well as positive control. Plates were washed four times with 0.05% PBST and blocked for 1 h with 200 μL/well PBS + 1% BSA. Plasma was diluted 1:50 in 0.05% PBST + 0.1% BSA and 50 μL was applied for 1.5 h. Plates were washed six times and total IgG binding was measured in a 1-h incubation with 100 μL/well of 1:50,000 HRP-conjugated polyclonal goat anti-human IgG (Jackson ImmunoResearch Labs) and developed using 50 μL/well TMB substrate (Sigma Aldrich). Reactions were stopped with an equal volume of 1 M H_2_SO_4_ and OD read on a BioTek synergy HT plate reader at 450 nm.

### Statistical analysis

Statistical analyses were performed using GraphPad Prism 9 with two-sided *P*-values < 0.05 considered significant. Normality of data distributions were assessed using Shapiro-Wilk test. Significance in correlations were assessed using Spearman’s rank correlation coefficient. Differences in means with standard deviation (SD) or medians with interquartile range (IQR, calculated as IQR = Q3 - Q1) between groups were compared by using one-way ANOVA, Student’s *t*-test, Friedman test or Mann-Whitney *U*-test as appropriate based on normality of data distribution.

## Author Contributions

KAH, RJS, CAF and KB designed and conducted the experiments, analyzed data, constructed figures and illustrations and wrote/edited the manuscript. MDG and EW co-wrote and edited the manuscript. KMH and DPI assisted with experiments. DH coordinated patient consent, questionnaires and collected blood samples. MDG, EW, RJS and KAH obtained funding.

## Acknowledgments

We thank all participants for providing samples, those who recruited them and the associated teams. This includes interviewers, nurses, computer and laboratory technicians, clerical workers, research scientists, volunteers, managers and receptionists. This work was supported by a COVID-19 rapid research funding opportunity grant (VR1 – 173202) from the Canadian Institutes for Health Research awarded through the COVID Immunity Task Force, and the MRC (MR/V028448/1, MR/S00971X/1). The UK Medical Research Council and Wellcome (Grant ref: 217065/Z/19/Z) and the University of Bristol provide core support for ALSPAC. The funders had no role in study design, data collection and analysis, decision to publish, or preparation of the manuscript. Vector pcDNA3.1(-) containing the SARS-Related Coronavirus 2, Wuhan-Hu-1 S Glycoprotein Gene, NR-52420 was contributed by David Veesler and vector pCMV/R containing the SARS-Related Coronavirus 2, S Glycoprotein Gene, Lineage B.1.1.529, Omicron Variant, NR-56470 was contributed by JR Mascola for distribution through BEI Resources, NIAID, NIH. SARS-Related Coronavirus 2 S Protein Peptide Array, NR-52402 was obtained through BEI Resources, NIAID, NIH. Adobe Illustrator was used to construct figures.

## Data Availability Statement

Data supporting the findings of this study and preserving anonymity of study participants will be made available from the corresponding authors through electronic correspondence for legitimate scientific purposes.

